# Incorrect Molecular Weights due to inaccurate Prestained Protein Molecular Weight Markers that are used for Gel Electrophoresis and Western Blotting

**DOI:** 10.1101/2020.04.03.023465

**Authors:** Rômulo Leão Silva Neris, Ajuni Kaur, Aldrin V. Gomes

## Abstract

The most widely used Western blotting protein standards are prestained proteins of known molecular mass (kDa). They are also utilized for sodium dodecyl sulphate (SDS) Polyacrylamide Gel Electrophoresis (PAGE) to determine the molecular mass of proteins separated by electrophoresis. The objective of this study was to assess the reliability of different commercially available protein standards in predicting accurate protein molecular weights. We performed this experiment by running Criterion TGX gels with five prestained protein standards (Thermo Fisher SeeBlue Plus 2, Bio-Rad Precision Plus Protein Dual-color, Thermo Fisher Spectra Multi-color, Novex-Sharp Pre-stained, and Invitrogen iBright Pre-Stained). To evaluate their accuracy, we utilized highly purified Bovine Serum Albumin (BSA, 66.44 kDa) and Cytochrome C (Cyto C, 11.62 kDa). We also made use of the dimers of BSA (132.88 kDa) and Cyt C (23.24 kDa) that are present on SDS-PAGE gels. Our results suggest that three of the standards were less accurate at higher molecular masses with the iBright marker having the highest error in determining the expected 132.88 kDa molecular weight. The SeeBlue Plus 2 was accurate at identifying the 132.88 kDa molecular weight protein band but was less reliable for the three other lower molecular weight proteins. These findings have significant implications for the determination of protein masses because researchers rely on these standards to evaluate the molecular masses of their protein(s). We suggest that at least two different protein standards should be initially used in electrophoresis gels and for Western blotting in order to get accurate protein molecular weight results.

## INTRODUCTION

Protein standards, also known as protein markers or protein ladders, were first developed in the early 1970’s (1). Over the last decade, the life science industry has created several types of protein molecular weight standards. These standards are typically composed of tagged purified proteins of known molecular masses to act as a reference protein standard. A typical tag is a chromophore bound to the protein, resulting in a prestained protein standard. SDS-PAGE, Western Blots and Isoelectric focusing are some of the techniques that rely on the use of protein markers as standards to determine the molecular weight of proteins of interest. This process can be easily performed because standards are distributed in a ready-to-use format and require no dilution, heating or addition of reducing agent. Once loaded into the gel-wells, prestained standards can be observed as the gel electrophoresis is being run. The molecular weights gained from the standards on gels can then be used to determine the molecular weight of the protein of interest.

Obtaining the correct molecular weight of a protein is critical for many experiments. For example, validating that an antibody recognizes a target protein usually involves the antibody recognizing a protein of a known molecular weight. Proteins that run at unexpected molecular weights sometimes contain a region rich in negatively charged amino acids or may have post-translational modifications such as glycosylation (2, 3).

A neighboring lab performed a Western blot and included prestained standards from two different commercial suppliers to determine which standard had brighter and easier to observe bands. They visited our lab when they observed that the two standards gave them different molecular weights for their target protein. They wanted to determine which standard was best to “trust.” They were told that they need to use purified proteins with well-established molecular weights to determine which is more accurate. They chose the standard that they have been using more often, so we decided to investigate how much variation in the molecular weights determined using different prestained standards currently exist.

We hypothesize that some molecular weight standards would not be as accurate as other standards. To investigate this hypothesis, we utilized SDS-PAGE to compare five different commercially available prestained molecular weight standards with purified proteins of well-established molecular weights. Using the different prestained molecular weight standards, we observed that some molecular weight standards were less accurate with respect to either low or high molecular weight determinations.

## METHODS

### Preparing samples

The standards were used as provided by the suppliers. The following protein standards were used: See Blue Plus (SB; Cat # LC5925; Lot 2064863 [first batch], Lot 2141778 [second batch]); Precision Plus Protein Dual Color Standards (PP; Cat No 161-0374; Lot L001648A); Spectra Multicolor Broad Range Protein Standard. (SM; Cat No 26634; Lot 00134245); Novex Sharp Prestained Protein Ladder (NS; Cat No 57318; Lot 2115570) and Invitrogen iBright Prestained Protein Ladder (iB; Cat No LC5605; Lot 00778110). The following purified proteins were also used to determine the accuracy of the standards: albumin (98-99% purity, Sigma-Aldrich, Inc, Cat. No A-3803; Lot 33H0665) and cytochrome C type III (97% purity, Sigma-Aldrich, Inc, Cat. No C-2506; Lot 45F-7195). These proteins were prepared by dilution in 10 mM Tris, 50 mM NaCl, pH 7.5 to a final concentration of 1 μg/μl. Protein concentrations were measured using the absorbance at 280nm on a Nanodrop 2000c spectrophotometer (Thermo Scientific, Inc). Samples were diluted 1:1 in SDS Sample Buffer (240 mM Tris, 8% [w/v] SDS, 40% [v/v] Glycerol, 0.4% [w/v] Bromophenol Blue, 5% [v/v] 2-Mercaptoethanol) and heated for 10 minutes at 95° C.

### Gel Electrophoresis

Experiments were run using 26-well gels (Criterion TGX Precast Gels, 15 μl, 1.0 mm, 4-20%, Bio-Rad Laboratories, Inc. Cat No 5671095). 13 wells were loaded by one researcher and the other set of 13 wells were loaded by another researcher for each set of experiments. This latter approach allowed us to determine the reliability of loading of the bands. The first five wells of each set of 13 wells were the commercial molecular weight standards, while the next seven consisted of the purified proteins. The amount (by volume) of the prestained standard to use for each lane was determined using test runs due to some standards having different concentrations of proteins than others (resulting in lighter bands than the others). For the protein standards, the following volumes were used: See Blue Plus (3 μl), Precision Plus Protein Dual Color Standards (1 μl), Spectra Multicolor Broad Range Protein Standard (8 μl), Novex Sharp Prestained Protein Ladder (4 μl) and Invitrogen iBright Prestained Protein Ladder (2.5 μl). The running time for all gels was 1hr 25min at 120V, which allowed for the samples to migrate through the gel. The gels were run in Tris-Glycine Buffer (25 mM Tris, 192 mM glycine, 0.1% [w/v] SDS) and the Bio-Rad PowerPac HC (Bio-Rad, Inc) was used as a power source. Following the running time, gels were carefully removed from the Bio-Rad Criterion Cell Container and then stained with Coomassie Blue R solution (0.025% [w/v] Coomassie R-250, 40% [v/v] Methanol, 7% [v/v] Acetic Acid). The gels were stained for an hour and then destained (Destaining Solution: 50% [v/v] Methanol, 7% [v/v] Acetic Acid) to remove non-specific background. Destaining was done in a two-step process: an hour on the shaker then another step in the destaining solution overnight without shaking.

### Gel Analysis

Gel scans and images were taken and imported to ImageJ Software (4). The distance from the middle of each band up to the beginning of the corresponding well was measured in pixels using the straight line tool, keeping an exact 90° inclination in the line. The lengths were stored using the region of interest (ROI) manager tool and then were exported to a Microsoft Excel Software spreadsheet. For each standard, we generated a scatter graph with the length (pixels) and Log (manufacturer’s predicted molecular weight) of the measured bands. The following step was to obtain a linear regression fit [f(x) = ax + b] with this data and interpolate the unknown lengths to obtain the relative molecular weight. Next, we calculated the error percentage by comparing the predicted protein molecular weights and the calculated protein molecular weight. According to the manufacturer, BSA had a size of 66.4 kDa. To compare their weights, both sequences were accessed on the Uniprot database (P02769 [ALBU_BOVIN] and P00004 [CYC_HORSE], respectively). Their molecular weights were predicted using Protein Molecular Weight prediction tool available at https://www.bioinformatics.org/sms/prot_mw.html. The blast sequence from the mature BSA form (in between positions 25-607) had a predicted size of 66.44 kDa. Equine Cytochrome C complete blast sequence had a predicted size of 11.62 kDa. Their dimers were considered twice this weight, with 133.88 and 23.24 kDa, respectively.

### Peer-reviewed publications analysis

To analyze how researchers are reporting the use of molecular weight markers, the US National Library of Medicine (PubMed, National Institutes of Health, available at https://www.ncbi.nlm.nih.gov/pubmed/) was accessed. A search for the term “Western Blot” was made and the results were filtered according to the following parameters: only papers with free full text available, published in January 2020 (01-31). The compiled searching parameters as presented in the webpage textbox “Search Details” was (“blotting, western”[MeSH Terms] or (“blotting”[All Fields] and “western”[All Fields]) or “western blotting”[All Fields] or (“western”[All Fields] and “blot”[All Fields]) or “western blot”[All Fields]) and (“loattrfree full text”[sb] and (“2020/01/01”[PDAT]: “2020/01/31”[PDAT])). This search was made in February 25^th^, 2020 and 877 papers were returned from these parameters. The following information was collected from each one: DOI number, do the authors use ladders in this publication, do they report ladders’ manufacturer and brand in methods or legends, do they show the ladder along with WB images, do they show the ladders’ equivalent band weight with WB images, do they show target protein expected size.

### Statistical analysis

One-way ANOVA or Two-way ANOVA followed by Dunnett’s multiple comparisons test were performed to analyze the data using GraphPad Prism version 7.00 for Windows, GraphPad Software, La Jolla California USA, www.graphpad.com.

## RESULTS

Standards with different colors on specific proteins separated as expected and appeared similar to the manufacturer’s images of the resolved standards on gels (Figure 1 and Supplemental Figure 1). The coomassie stained gel showing different commercial prestained standards are labeled according to the manufacturer’s information is shown in Figure 1A.

**Figure 1.**
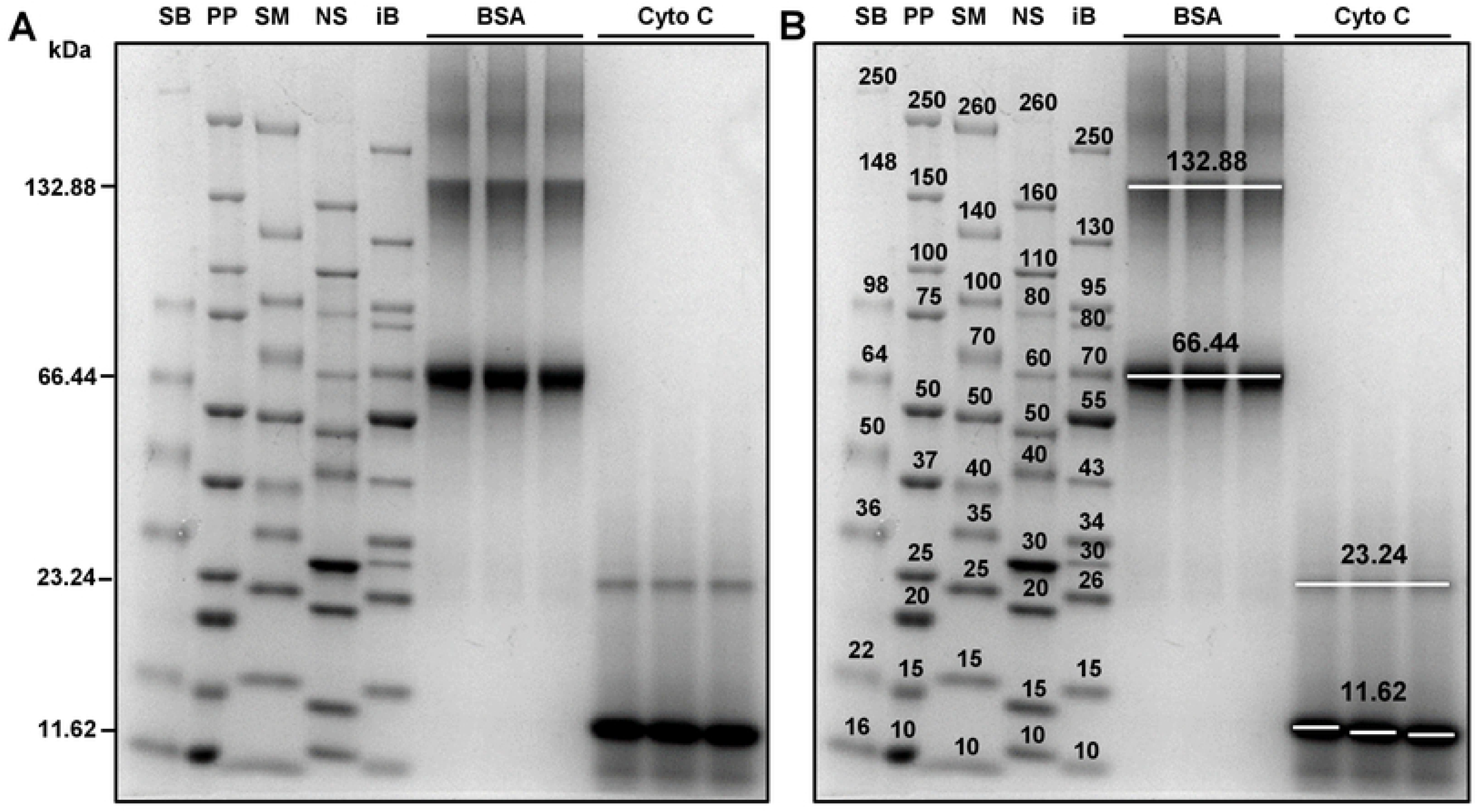
Gel Electrophoresis Protein standards exhibit relative mobility discrepancies. Five different prestained protein ladders (SeeBlue Plus 2 (SB), Precision Plus Dual (PP), Spectra Multicolor (SM), Novex Sharp (NS), and iBright (iB)) were run along with two different purified proteins (bovine Serum Albumin (BSA) and Cytochrome C (Cyto C)) in a Tris/Glycine 4-20% pre-cast gel. (A) Representative image of a Coomassie-blue R stained-gel. The molecular weights on the left side of the gel are based upon BSA and Cyto C. (B) Representative image of the same Coomassie-blue R stained-gel in (A) with each band labeled by size in kilodaltons (kDa). White lines mark the region where the mobility was measured. Bands were labelled according to the manufacturer’s size recommendations. The image is representative of six experiments performed by two different researchers.

### Inconsistencies in Molecular Weights of Commercial Protein Standards

Unexpectedly, each of the five protein standards resolved differently with detectable discrepancies in the molecular weights of the proteins that make up the different standards (Figure 1B). The 22 kDa band for the SeeBlue Plus 2 standard was aligned with the 15 kDa band of Spectra Multicolor standard (Figure 1B). The 15 kDa band of the Spectra Multicolor standard was not aligned with two other 15 kDa bands from other standards. Similar discrepancies were observed around the 50 kDa region, the 100 kDa region and the 250 kDa region (Figure 1B). The 100 kDa band of the Precision Plus Protein Dual Color Standard had the same mobility as the 110 kDa band of the Sharp Prestained Protein Ladder.

### Protein Molecular Weight Determination

In order to evaluate the inaccuracy of these protein ladders in gel electrophoresis molecular weight determinations, two different proteins were chosen for comparison: BSA and horse Cyto C type III. These proteins’ molecular weights and sequences are well known and studied in the last decades (1). They were also chosen because they can form stable oligomers like dimers and trimers in solution in many different conditions. Cyto C is able to form dimers by swapping the C-terminal domains in aqueous solutions (5). Aged albumin is able to aggregate by the oxidation of two-free-SH groups or by changing of a single -SH group with a S-S bridge in another monomer (6). Non-aged albumin is also able to self-associate and form oligomers, mainly as dimers and in a reversible way (7). BSA and Cyto C consistently appear both as monomers and dimers by electrophoresis as described previously (1), and these forms can be used to access the efficiency of different ladders. These two proteins and their dimers allowed us to determine a range of molecular weights on the polyacrylamide gel (Cyto C, 11.62 kDa; Cyto C dimer, 23.24 kDa; BSA, 66.44 kDa; BSA dimer, 132.88 kDa) (Figure 1B).

For each standard, a linear regression fit was performed to determine the molecular weights of BSA and Cyto C (Figure 2). The linear regression fits were of relative mobility of the highest molecular weight to the lowest molecular weight band of each standard. The R^2^ value was accessed and values greater than 0.97 were found for all linear regressions (Data shown in Figure 2 legends).

**Figure 2.**
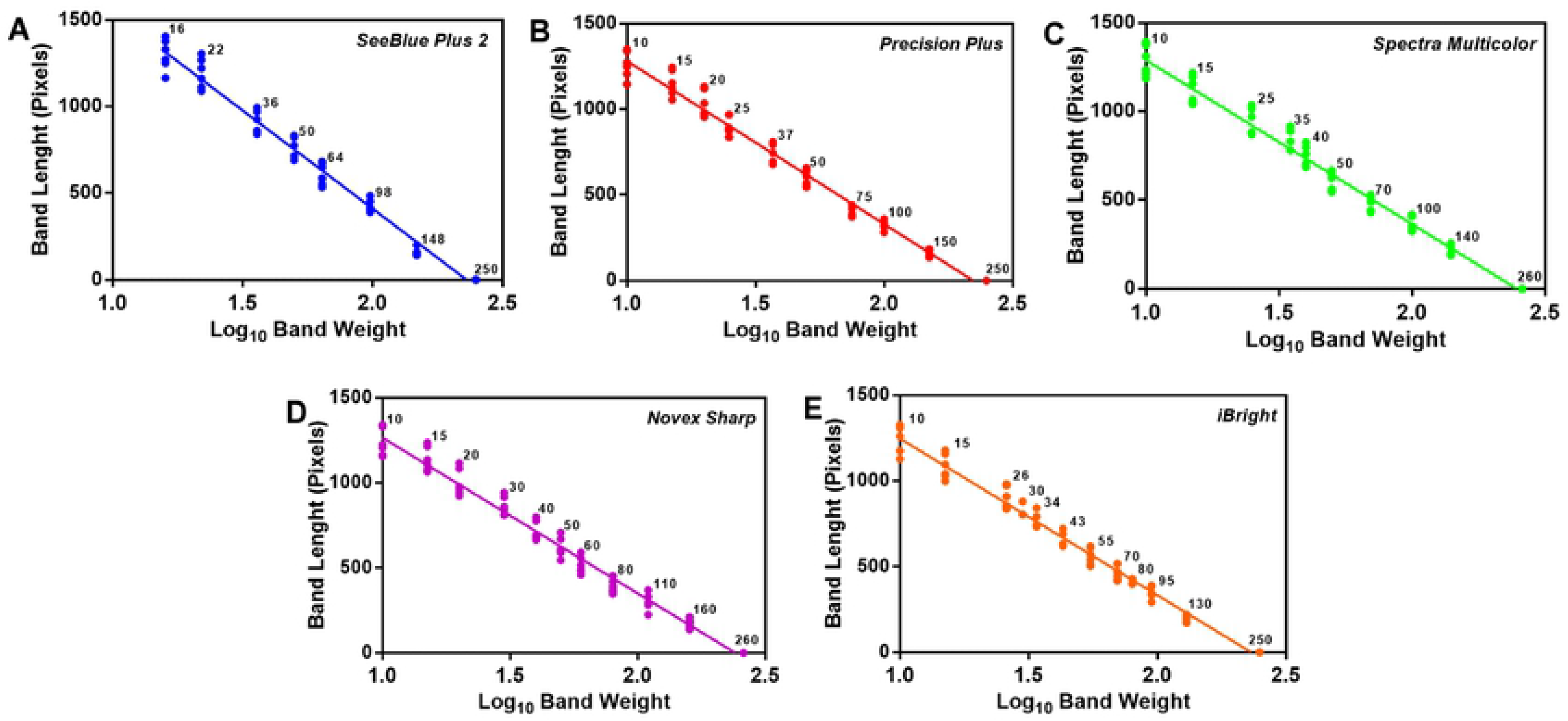
Linear regression of protein size and relative mobility for each protein standard. Five different protein ladders were run in a Tris/Glycine 4-20% pre-cast gel, and their mobility fit to the protein molecular weight that was given by the manufacturer. (A) SeeBlue Plus 2 (R^2^ = 0.979), (B) Precision Plus Dual (R^2^ = 0.9808), (C) Spectra Multicolor (R^2^ = 0.9773), (D) Novex Sharp (R^2^ = 0.9773), (E) iBright (R^2^ = 0.9803). The number next to each set of dots is the manufacturer expected protein size in kDa. Simple linear regression, equation and R^2^ was calculated for all data sets. The results represent six experiments performed by two researchers.

### Differences in Molecular Weight of Cytochrome C as calculated using different Commercial Protein Standards

Each protein standard performed differently when compared to the BSA and Cyto C (Table 1 and Figure 3). Only SeeBlue performed poorly with respect to determining the molecular weight of Cyto C monomer. The average molecular weight of Cyto C (11.62 kDa expected) predicted by the SeeBlue standards was 22.12 ± 5.38 kDa which is about 90% higher than the expected molecular weight (Figure 3A). The standard with the closest predicted molecular weight to the expected molecular weight was the Precision Plus Dual standard with a predicted molecular weight of 12.32 ± 1.44 kDa. The other standards showed 15.77% to 21.43% higher predicted molecular weight for the Cyto C monomer. The predicted molecular weight of the Cyto C monomer was significantly different between the Seeblue Plus standard and each of the other four standards (P < 0.001).

**Table 1.** Predicted molecular weights of cytochrome C and bovine serum albumin using different commercial pre-stained standards.

| Protein | Predicted Weight | See Blue Plus 2 (Lot # 2064863) | Precision Plus Dual | Spectra-Multicolor | Novex Sharp | iBright |
| --- | --- | --- | --- | --- | --- | --- |
| Cyto C | 11.62 kDa | 22.12 ± 5.38<sup>†</sup><br>(90.33 %) | 12.32 ± 1.44<br>(6.01 %) | 13.45 ± 0.74<br>(15.77 %) | 13.58 ± 1.22<br>(16.86 %) | 14.11 ± 0.73<sup>†</sup><br>(21.43 %) |
| Cyto C Dimer | 23.24 kDa | 35.69 ± 5.96<sup>†</sup><br>(53.55 P%) | 23.41 ± 2.49<br>(0.73 %) | 25.85 ± 1.95<br>(11.22 %) | 26.21 ± 2.21<br>(12.80 %) | 21.17 ± 1.56<br>(-16.92 %) |
| BSA | 66.44 kDa | 73.88 ± 7.38<br>(11.20 %) | 61.62 ± 3.28<br>(-7.25 %) | 69.21 ± 4.73<br>(4.17 %) | 66.91 ± 2.08<br>(0.71 %) | 73.21 ± 2.34<br>(10.19 %) |
| BSA Dimer | 132.88 kDa | 138.22 ± 10.71<br>(4.02 %) | 142.51 ± 6.93<br>(7.25 %) | 162.19 ± 13.96<sup>†</sup><br>(22.05 %) | 156.64 ± 6.21<br>(17.88 %) | 171.44 ± 8.10<sup>†</sup><br>(29.02 %) |

Values are mean ± SD (n=6 for all values). Values in brackets are the % difference in predicted vs expected values. <sup>†</sup> Predicted protein molecular weights that are > 20% the expected molecular weight.

**Figure 3.**
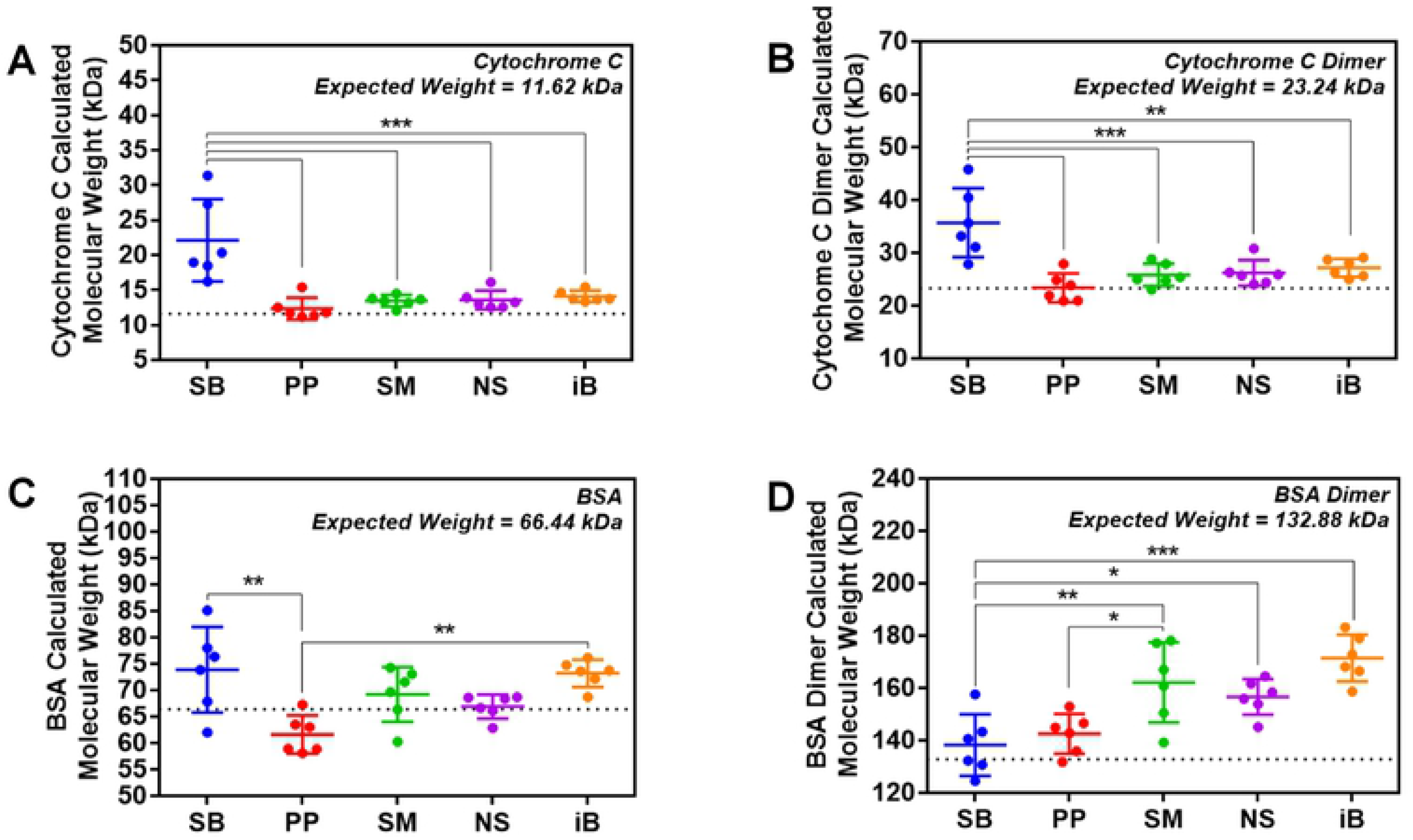
Bovine Albumin and Cytochrome C molecular weight differences according to each protein standard. Two purified proteins with well-established molecular weights (BSA and Cyto C) were run with the five different protein standards on a Tris/Glycine 4-20% pre-cast gel and the molecular weights of BSA and Cyto C calculated based upon the molecular weights of the standards obtained from the manufacturers of the standards. (A) Calculated molecular weight of Cytochrome C; (B) Calculated molecular weight of Cytochrome C dimer; (E) Calculated molecular weight of BSA; (D) Calculated molecular weight of BSA dimer. Dot lines (···) represent the expected molecular weight for each protein. Data are shown as mean ± S.D. n = 6 experiments performed by two researchers. *p<0.05; **p<0.01.

The molecular weight prediction of the Cyto C dimer (23.24 kDa expected molecular weight) was also least accurate when the SeeBlue Plus was utilized (35.69 ± 5.96 kDa, about 53% higher than the expected molecular weight) and most accurate when using the Precision Plus Dual (23.41 ± 2.49 kDa). The other three standards also showed moderate accuracy with predicted molecular weights of 21.17 to 26.21 kDa (Figure 3B). The predicted molecular weight of the Cyto C dimer was significantly different between the Seeblue Plus standard and each of the other four standards (P < 0.01).

### Differences in Molecular Weight of Bovine Serum Albumin as calculated using different Commercial Protein Standards

The BSA monomer (66.44 kDa) molecular weight prediction using the Novex Sharp standard was the most accurate (66.91 ± 2.08 kDa) (Table 1 and Figure 3C). The Seeblue Plus standard had much better accuracy in predicting the BSA monomer’s molecular weight (73.88 ± 7.38kDa, 11.20% higher than expected) than the Cyto C molecular weights. The other three standards had predicted molecular weights varying from 61.62 ± 3.28 kDa (Precision Plus Dual) to 73.21 ± 2.34 kDa (iBright) (Figure 3C). The molecular weights predicted by SeeBlue Plus and Precision Plus Dual as well as the molecular weights predicted by Precision Plus Dual and the iBright standards were statistically significant (P < 0.01).

Only two of the standards were accurate (less than 10% difference from expected molecular weight) at determining the molecular weight of the BSA dimer (132.88 kDa). The most accurate standards for predicting the molecular weight of the BSA dimer were SeeBlue Plus (138.22 ± 10.71 kDa) and Precision Plus Dual (142.51 ± 6.93 kDa). Spectra Multicolor and Novex Sharp standards had higher predicted molecular weights of 22.05 and 17.88% and the iBright standard had the largest variation from the expected molecular weight and was 29% higher than expected (Figure 3d).

### Lot to Lot variation in the SeeBlue Plus 2 molecular weight standard

Since SeeBlue Plus 2 had the largest differences in predicted and expected low and medium molecular weight determinations, a second lot (batch) of SeeBlue Plus 2 was purchased and investigated to determine if the deviation from expected molecular weights was intrinsic to the molecular weights assigned to the prestained molecular weights or if it was lot-specific. Analysis of a second lot of SeeBlue Plus 2 showed that this lot exhibited the same range of differences in comparison with other standards (Supplemental Table 1). The molecular weight regions corresponding to 10 kDa, 50 kDa and 100 kDa show apparent differences with other standards (Supplemental Figure 1). The linear regression curves for both lots of SeeBlue Plus were similar, with no statistically significant differences between them (Supplemental Figure 2).

This SeeBlue Plus lot#2 exhibited significant differences in Cyto C monomer and its dimer molecular weight prediction when compared to the other four ladders, since to the first lot of SeeBlue Plus standard used (Supplemental Figure 3). The SeeBlue Plus lot#2 showed higher predicted molecular weights of 43.27% and 38.62%, for the Cyto C monomer and dimer respectively. In BSA and BSA dimer prediction size, no significant differences were found among the tested standards. However, all five standards had errors of more than 15% for BSA dimer molecular weight prediction (Supplemental Table 1).

### Reporting of molecular weight standards and predicted molecular weights of target proteins in the literature

Next, we investigated if information regarding the use of standards was being reported in the literature. We searched the PubMed database for free full access papers published in January 2020 that used western blotting. This search returned 877 papers, that were examined to profile how standard information is reported. We examined how standards technical information was being reported and how standard molecular weight data were shown in these papers. From the 877 papers investigated, 830 (94.64% of those examined) had no information about if standards were used, possibly because the authors did not use standards or because they use standards but do not mention them in the publications (Figure 4). 34 papers (3.88%) report the use of standards but lacked relevant information about the type or manufacturer of the standard. Only 13 articles (1.48%) used standards and reported the appropriate information about these standards. When these 877 papers were investigated to determine how they report target molecular weight data, 632 papers (72.06%) only showed target protein bands without any information regarding the molecular weights determined for the target protein. 192 (21.89%) publications reported the target molecular weight on the western blot image, without any information about if this molecular weight is the expected molecular weight or the molecular weight predicted from protein molecular weight standards. Only 53 papers (6.04%) showed protein standards (either partially or fully) together with the western blot.

**Figure 4.**
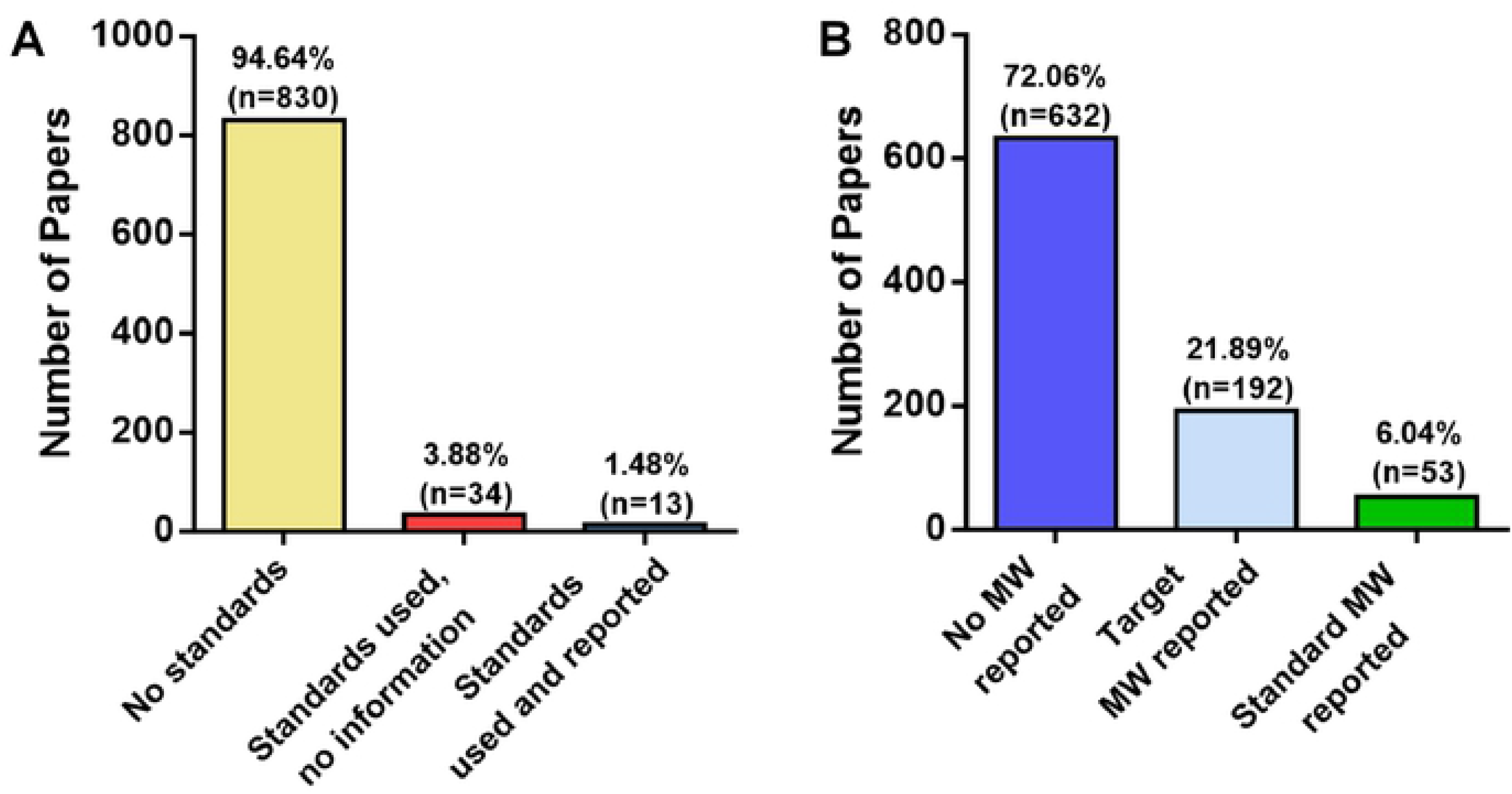
Literature analysis of protein standards usage and molecular weight reports. All papers on the PubMed database published in January 2020 for the search term “Western Blot” were accessed and manually investigated. In total, 877 publications were investigated. (A) Western blotting publications that used standards as well as the number of publications that reported the type of standard that was utilized. (B) Western blotting publications that report the molecular weight of the protein of interest as well as publications that show the molecular weights of the protein standards.

## DISCUSSION

Western Blotting is one of the most used techniques in life sciences to study protein amounts, distribution and modifications, such as phosphorylation. Protein molecular weight standards are widely used as references in these experiments to determine if the antibody is recognizing a target protein of the appropriate molecular weight. However, very few investigations on the reliability of protein standards have been carried out, and most of them were done several decades ago, before the use of commercial standards (1, 8, 9). It should be noted that proteins can show different molecular weights on different polyacrylamide gel concentrations so the results presented in this manuscript may be different when other polyacrylamide gels are used. Dunker et al. (1) showed that Cyto C and BSA intrinsic apparent weight variation on different polyacrylamide gel concentrations is lower than 9.7 and 6.5%, respectively. There are many different aspects that can lead to variations in protein electromobility and thus change the resolution of a protein band that is used as part of the standard. Many prestained molecular weight markers use dyes like remazol brilliant, which “stains” proteins by covalently binding its vinyl groups to amines, alcohol and sulfhydryl present in the proteins (10). Despite its ease of use, covalently bound remazol could decrease the electromobility of proteins in a non-linear way, showing different resolving distances during electrophoresis in comparison with unstained proteins (11). Similar labeling of proteins can be achieved using other dyes with the same principle, like drimarene blue and dabsyl chloride (12, 13).

Having relatively accurate molecular weight determinations are important for gel electrophoresis and western blotting because many antibodies cross-react with other non-target proteins resulting in false positives (14, 15). Additionally, knowing the molecular weight of bands detected by western blotting is one of several techniques to ensure the antibody is detecting the expected molecular weight protein. When the molecular weight is different from what is expected that molecular weight is also important because it is usually associated with post-translational modifications of the target protein. Protein modifications like phosphorylation can increase the molecular weight of a protein during electrophoresis (16). Glycosylated protein content when heterogeneously present among the proteins can be responsible for high variations in protein mobility through electrophoresis. Wang and collaborators showed that the non-glycosylated protein sCD38 has a theoretical size of 30.5 kDa on a polyacrylamide gel. Glycosylation of sCD38 increased its observed size by 25.5%, giving it a predicted molecular weight of 37.1-39.5 kDa (17). Another factor that can interfere with migration and resolution are increased salt concentrations in the buffers used for protein preparation for electrophoresis (18). Many different proteins have slightly different migration patterns with the increase of salt concentration in the electrophoresis buffers, including albumin, ovalbumin, lactate dehydrogenase, and trypsin inhibitor. Under certain mechanical or chemical adverse conditions, like when metallic ions are in abundance, some proteins can undergo proteolysis, and variations can appear according to the amount of protein and the part of the target protein cleaved during the process (19). Specific proteins, like membrane proteins, are also known to show anomalously migration during electrophoresis (20). However, a quick review of the literature suggests that >90% of target proteins (even those that are post-translationally modified) run at their predicted molecular weight (data not shown).

Our data suggest that protein standards present more substantial variations than expected when used under the exact conditions. These findings raise a special concern about how prestained protein standards molecular weights are calibrated and how the use of standards and molecular weight determinations are reported in the literature. Inaccurate standards could lead to misinterpretations and errors in the analysis due to significant variations in the calculated target protein weight. Two different researchers using the same conditions and samples but using different protein standards to investigate the molecular weight of the same target protein may get significantly different results.

Among the five standards analyzed in this paper, SeeBlue Plus 2 was the least accurate with small proteins molecular weight predictions, with calculated molecular weights that were 90.33% and 53.55% higher than expected for Cyto C and its dimeric form, respectively. The Precision Plus was accurate at predicting the protein molecular weights of all the bands investigated. Monomeric Cyto C, dimeric Cyt C, monomeric BSA and dimeric BSA calculated molecular weights were less than 8% altered from the expected molecular weights. Both Spectra Multicolor and Novex Sharp ladders had similar profiles, with differences from the expected molecular weights ranging from 13.45% and 13.58% for the prediction of Cyto C monomer and 11.22% and 12.80% for the Cyto C dimer. The Spectra Multicolor and Novex Sharp standards had less than 5% differences from the expected molecular weights for monomeric BSA, and 22.05% and 17.88% for dimeric BSA, respectively. The iBright standards had variations from 10.19% to 21.43% for Cyto C monomer, Cyto C dimer and BSA monomer. The iBright standards showed the most significant difference in the expected molecular weight for the BSA dimer with a calculated molecular weight that was 29% higher than expected. Interestingly, the iBright standard is a mixture of twelve recombinant proteins, with ten prestained proteins and two proteins (molecular weights 30 kDa and 80 kDa) that are unstained. Unstained standards would likely be more accurate than prestained standards since molecular weights of unstained standards are better documented and may account for why the iBright standard is better at predicting molecular weights near the unstained standards.

These variations were found by two independent researchers with different backgrounds (one a 4^th^ year undergraduate and the other a 5^th^ year graduate student), suggesting that these variations are very likely to be inherent for the use of standards. Support for the suggestion that the difference in expected and predicted molecular weights are due to molecular weights assigned to the pre-stained standard protein bands comes from similar results being obtained when a different lot of SeeBlue Plus 2 standards was used. In both batches of standards, significant errors in the calculated molecular weights of monomeric Cyto C and dimeric Cyto C were obtained relative to the expected molecular weights. Hence, the variations observed are not only associated with human-manipulation errors, but intrinsic variations that can be carried to multiple experiments and generate incorrect data analysis for protein molecular weights. The higher error related to calculating the molecular weight of the BSA dimer is likely due to changes in the mobility of the standards with different batches of the SDS-PAGE gel since all other standards showed higher errors than in the first set of experiments. It is important to note that most prestained manufacturers state that their standards will allow you to determine the approximate or apparent molecular weight of the protein of interest. ThermoFisher Scientific recommends using unstained protein ladders for molecular weight determinations. However, of the 877 publications investigated, 6.04 % of manuscripts reported the standards they used were not unstained standards.

The concerns associated with incorrect molecular weight determinations is compounded by the lack of reporting of how molecular weights of target proteins are carried out, and if they are even determined in some cases. The ability to replicate a study that used molecular weight standards to determine the molecular weight of a target protein by western blotting may need to use the same molecular weight standards. However, only about 1.5% of manuscripts (out of 830) report what molecular weight standards they are using. While some authors show the standards and the molecular weights of these standards, they do not state the name and source of these standards. Disappointingly, only approximately 22% of publications investigated reported molecular weights for the target bands they identified. This lack of reporting makes it difficult to uncover possible inaccuracies related to the protein standard used.

## CONCLUSION

Overall, our results suggest that researchers need to be aware that different prestained molecular weight standards can result in different molecular weights of proteins being calculated. Researchers should note that some standards were accurate for certain, but not all molecular weight ranges. Researchers could run a purified protein with a well-established molecular weight close to their target protein to validate that the protein standards they are using are accurate for that molecular weight range of interest. Another possibility is to use more than one standard at a time. Using an unstained protein standard that can be stained (with Ponceau S for example) on the western blotting membrane (before blocking or after the western blotting) together with a prestained standard could also help to validate the prestained standard which can then be used by itself for subsequent western blots. Researchers should report what protein standards they are using and label molecular weights on western blots used in publications to increase the reproducibility of this method.

## Acknowledgements

This research was supported by grants from the National Institutes of Health Superfund Research Program (P42 ES004699) and the American Heart Association (16GRNT31350040).

## SUPPORTING INFORMATION CAPTIONS

**S1 Text. Protein standards exhibit relative mobility discrepancies.** Five different prestained protein ladders (SeeBlue Plus 2 (SB), Precision Plus Dual (PP), Spectra Multicolor (SM), Novex Sharp (NS), and iBright (iB)) were run along with two different purified proteins (ovine Serum Albumin (BSA) and Cytochrome C (Cyto C)) in a Tris/Glycine 4-20% pre-cast gel. (A) Representative color image of a Coomassie-blue R stained-gel. The molecular weights on the left side of the gel are based upon BSA and Cyto C. (B) Representative color image of the same Coomassie-blue R stained-gel in (A) with each band labeled by size in kilodaltons (kDa). White lines mark the region where the mobility was measured. Bands were labeled according to the manufacturer’s size recommendations. The image is representative of six experiments performed by two different researchers.

**S2 Text. Linear regression between protein molecular weight and relative mobility for two different lots (batches) of SeeBlue Plus 2.** The linear regression fit of two different lots of SeeBlue Plus 2 significantly overlapped. The number next to each set of circles is the manufacturer expected protein size in kDa. The simple linear regression, equation and R^2^ were calculated for all data sets. n = 4 (SB batch #2) or 6 (SB batch #1).

**S3 Text. Bovine Albumin and Cytochrome C calculated size differences according to a second lot of SeeBlue Plus 2.** Two purified proteins with well-established molecular weights (BSA and Cyto C) were run with the five different protein standards on a Tris/Glycine 4-20% pre-cast gel, and the molecular weights of BSA and Cyto C calculated based upon the molecular weights of the standards obtained from the manufacturers of the standards. (A) Calculated molecular weight of Cytochrome C; (B) Calculated molecular weight of Cytochrome C dimer; (E) Calculated molecular weight of BSA; (D) Calculated molecular weight of BSA dimer. Dotted lines (···) represent the expected molecular weight for each protein. Data are shown as means ± S.D. n = six experiments performed by two researchers. *p<0.05; **p<0.01

## REFERENCES

1. Dunker AK, Rueckert RR. Observations on molecular weight determinations on polyacrylamide gel. The Journal of biological chemistry. 1969;244(18):5074–80.

2. Wilson N, Simpson R, Cooper-Liddell C. Introductory glycosylation analysis using SDS-PAGE and peptide mass fingerprinting. Methods in molecular biology. 2009;534:205–12.

3. Shi Y, Mowery RA, Ashley J, Hentz M, Ramirez AJ, Bilgicer B, et al. Abnormal SDS-PAGE migration of cytosolic proteins can identify domains and mechanisms that control surfactant binding. Protein science: a publication of the Protein Society. 2012;21(8):1197–209.

4. Schneider CA, Rasband WS, Eliceiri KW. NIH Image to ImageJ: 25 years of image analysis. Nature methods. 2012;9(7):671–5.

5. Hirota S, Hattori Y, Nagao S, Taketa M, Komori H, Kamikubo H, et al. Cytochrome c polymerization by successive domain swapping at the C-terminal helix. Proceedings of the National Academy of Sciences of the United States of America. 2010;107(29):12854–9.

6. Giancola C, De Sena C, Fessas D, Graziano G, Barone G. DSC studies on bovine serum albumin denaturation. Effects of ionic strength and SDS concentration. International journal of biological macromolecules. 1997;20(3):193–204.

7. Levi V, Gonzalez Flecha FL. Reversible fast-dimerization of bovine serum albumin detected by fluorescence resonance energy transfer. Biochimica et biophysica acta. 2002;1599(1-2):141–8.

8. Shapiro AL, Vinuela E, Maizel JV, Jr. Molecular weight estimation of polypeptide chains by electrophoresis in SDS-polyacrylamide gels. Biochemical and biophysical research communications. 1967;28(5):815–20.

9. Shapiro AL, Maizel JV, Jr. Molecular weight estimation of polypeptides by SDS-polyacrylamide gel electrophoresis: further data concerning resolving power and general considerations. Analytical biochemistry. 1969;29(3):505–14.

10. Compton MM, Lapp SA, Pedemonte R. Generation of multicolored, prestained molecular weight markers for gel electrophoresis. Electrophoresis. 2002;23(19):3262–5.

11. Saoji AM, Jad CY, Kelkar SS. Remazol Brilliant Blue as a pre-stain for the immediate visualization of human serum proteins on polyacrylamide gel disc electrophoresis. Clinical chemistry. 1983;29(1):42–4.

12. Bosshard HF, Datyner A. The use of a new reactive dye for quantitation of prestained proteins on polyacrylamide gels. Analytical biochemistry. 1977;82(2):327–33.

13. Parkinson D, Redshaw JD. Visible labeling of proteins for polyacrylamide gel electrophoresis with dabsyl chloride. Analytical biochemistry. 1984;141(1):121–6.

14. Gilda JE, Ghosh R, Cheah JX, West TM, Bodine SC, Gomes AV. Western Blotting Inaccuracies with Unverified Antibodies: Need for a Western Blotting Minimal Reporting Standard (WBMRS). PloS one. 2015;10(8):e0135392.

15. Ghosh R, Gilda JE, Gomes AV. The necessity of and strategies for improving confidence in the accuracy of western blots. Expert review of proteomics. 2014;11(5):549–60.

16. Lee CR, Park YH, Min H, Kim YR, Seok YJ. Determination of protein phosphorylation by polyacrylamide gel electrophoresis. Journal of microbiology. 2019;57(2):93–100.

17. Wang G, de Jong RN, van den Bremer ETJ, Parren P, Heck AJR. Enhancing Accuracy in Molecular Weight Determination of Highly Heterogeneously Glycosylated Proteins by Native Tandem Mass Spectrometry. Analytical chemistry. 2017;89(9):4793–7.

18. Seemann JR, Critchley C. Effects of salt stress on the growth, ion content, stomatal behaviour and photosynthetic capacity of a salt-sensitive species, Phaseolus vulgaris L. Planta. 1985;164(2):151–62.

19. Lyons B, Kwan AH, Truscott RJ. Spontaneous cleavage of proteins at serine and threonine is facilitated by zinc. Aging cell. 2016;15(2):237–44.

20. Rath A, Deber CM. Correction factors for membrane protein molecular weight readouts on sodium dodecyl sulfate-polyacrylamide gel electrophoresis. Analytical biochemistry. 2013;434(1):67–72.

